# In situ impedance measurements on postmortem porcine brain

**DOI:** 10.1101/2020.04.17.029728

**Authors:** Lucas Poßner, Matthias Laukner, Florian Wilhelmy, Dirk Lindner, Uwe Pliquett, Bojana Petkovic, Marek Ziolkowski, Thomas R. Knösche, Konstantin Weise

## Abstract

The paper presents an experimental study where the distinctness of grey and white matter of an in situ postmortem porcine brain by impedance measurements is investigated. Experimental conditions that would allow to conduct the same experiment on in vivo human brain tissue are replicated.

https://doi.org/10.1515/cdbme-2019-XXXX

## 1 Introduction

Knowing the exact electrical properties of brain tissues in individual subjects is important for modeling non-invasive brain measurement and stimulation techniques, such as EEG, MEG, and TMS. Moreover, such data may be important for the identification of pathological tissue, such as tumors. Such measurements are very challenging because (i) they are performed in living tissue at room temperature; (ii) due to the highly dispersive properties of brain tissue; (iii) a statistically relevant number of data sets need to be collected to derive statistically relevant parameters.

Previous studies were often based on inadequate and error-prone measurement procedures, such as monopolar measurements. Moreover, the studies were only conducted on a very small number of patients (N≈10) and for one particular frequency [1].Furthermore, substantial variability between experimental conditions, subjects and even repetitions of the same experiment could be observed, which can be caused by a number of influencing factors, including the day-dependent cognitive state, the basic hormone level or the current fluid balance. Since these factors are not known, the interpretation of the already small data sets is further complicated [2–4].

In order to overcome these drawbacks, we plan to record broadband conductivity spectra in living human brain tissue. The experiments are based on direct impedance measurement and will be performed on the occasion of brain tumor surgeries. A neurostimulator generates a periodic monophasic current pulse signal which is applied to the tissue via a certified stimulation electrode. The total current and the pulse width is variable. Both applied current and voltage across the electrode and the tissue are sampled and used to compute the impedance of the entire system. After the elimination of the electrode impedance the electric material properties can be computed using numerical electric field simulations.

Here, we report a proof-of-principle study, testing the feasibility of our measurement technique in a postmortem porcine brain. Our goal was to prove the distinctness of grey and white matter by their impedances.

## 2 Methods

### 2.1 Measurement preparation

The measurements were executed on postmortem porcine brain at its in situ position inside the head. The pig was regularly slaughtered in a slaughterhouse, using electric stunning. It was not slaughtered to conduct this experiment. Its head was separated from the body and the brain remained mechanically undamaged. By the time of the measurements it was dead for less than 12 hours.

Since it is permanently used in surgeries, it was not possible to use the neurostimulator for the actual experiment. To overcome this issue, we recorded stimulation pulses beforehand with an oscilloscope and used an arbitrary waveform generator to replicate them. We varied the current in the range of 10 µA to 100 µA and the pulse width in the range of 60 µs to 200 µs.

### 2.2 Measurement setup

For the actual measurement we used the setup shown in Fig.1.

**Figure 1:**
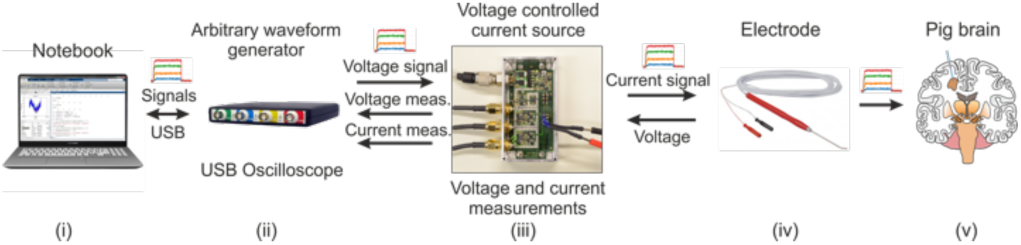
Measurement setup containing the laptop (i), the USB oscilloscope (ii), the electric circuits (iii), the brain stimulation electrode (iv) and the porcine brain (v) as the measurement object.

It consists of:

i. a laptop computer to control the signal generation and data acquisition;
ii. an USB oscilloscope (TiePie Handyscope HS5-540) with an integrated arbitrary waveform generator. The waveforms recorded from the neurostimulator were send to the arbitrary waveform generator and the oscilloscope was used to pick up the measurement signals from (iii);
iii. a measurement front end containing:
  - a voltage controlled current source that applies the stimulation pulse to the electrode and the tissue,
  - a differential amplifier that measures the voltage across the electrode and the tissue,
  - a differential amplifier that measures the stimulation current as a voltage drop across a sensing resistor [8];
iv. a bipolar coaxial brain stimulation electrode (Inomed BCS 45mm 30°) connected to the voltage controlled current source.

The electrode is an open-ended coaxial probe with an uninsulated outer conductor. Its end has a 30° bend and is 45 mm long. The cable length is 3 m. Due to its length, it yields an unwanted parasitic capacitance. This phenomenon occurs if a conductor is just slightly noncircular [5]. When compensating for the electrode impedance, this needs to be considered. However, in the context of this paper, presenting a proof of principle, this is negligible.

### 2.3 Measurement signals

It is of crucial importance to influence the brain tissue as little as possible. Hence, our goal was to identify the lowest possible current strength and pulse width that still allow to distinguish grey and white matter. As we have seen in prior experiments with plant tissues, the measured impedance strongly depends on the contact pressure between the electrode and the tissue as well as the humidity of the tissue surface. To eliminate the influence of these factors, we applied different combinations of current strengths and pulse widths in a short time span of about 0.8 s.

Therefore, we synthesized a signal containing all combinations of current strengths (10, 20, …, 100 µA) and pulse widths (60, 100, 200 µs). Each pulse was repeated 5 times with a duty cycle of 1/10. To limit the amount of charge at the electrode surface and therefore the DC surface potential we alternated the polarity of the current pulses [6]. The pulse train and an enlarged section of it are shown in Fig. 2.

**Figure 2:**
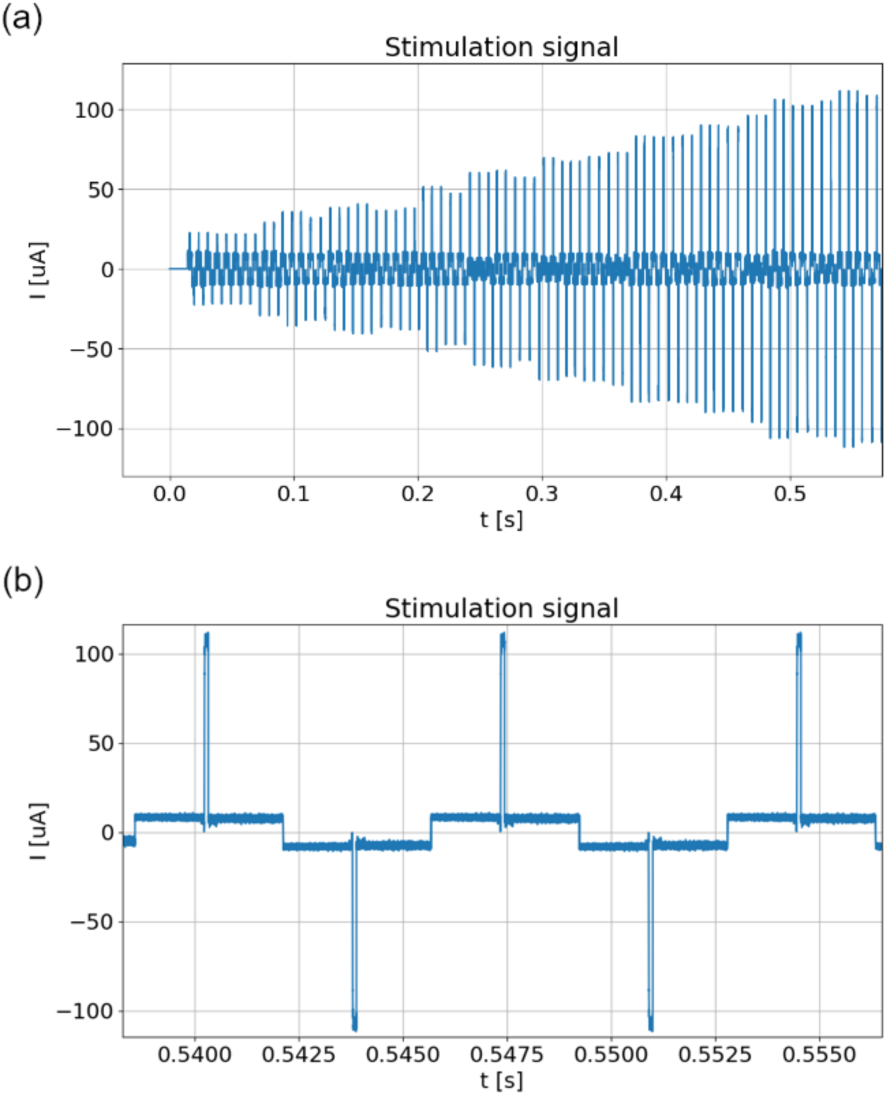
(a) Stimulation current pulse train containing 5 biphasic repetitions of each combination of current strength and pulse width; (b) Enlarged section of the current pulse train with 5 pulses.

### 2.4 Measurement procedure

A craniotomy was performed by a neurosurgeon on the pig head with neurosurgical instruments (Fig. 3a). The burr hole had a diameter of approximately 2 cm and the meninges were fixated with sutures giving access to the cortex (Fig. 3b).

**Figure 3:**
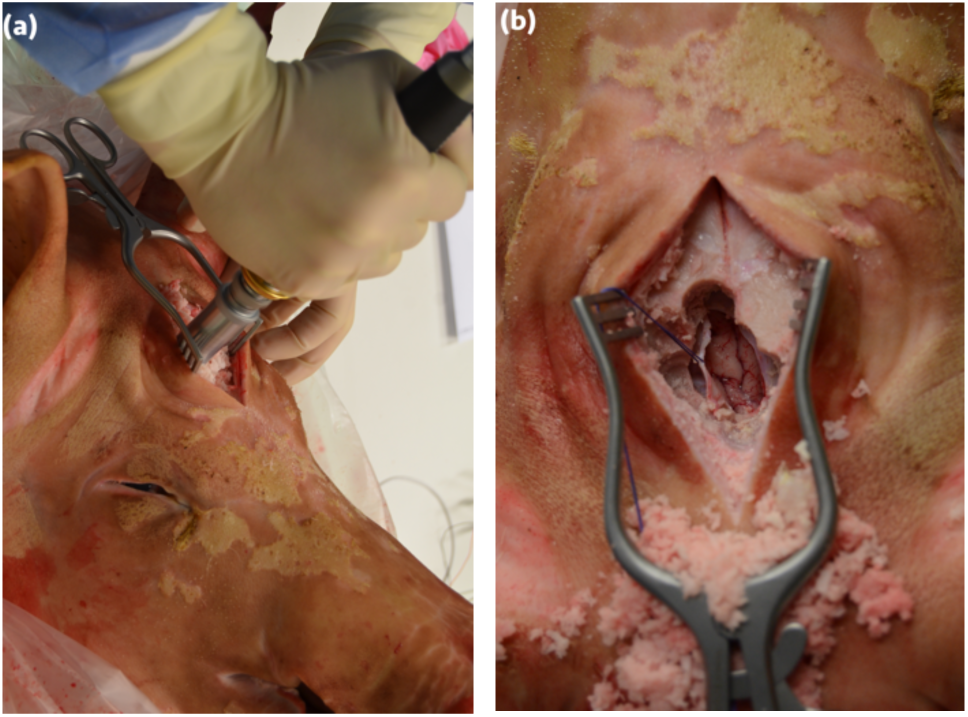
(a) Craniotomy on the pig head; (b) Burr hole with fixed meninges and access to cortex.

The cortex was cut with a scalpel in order to make grey and white matter accessible. The electrode was put on the tissue by hand with no additional fixation. Then the current signal was applied and both the stimulation current and the voltage response were recorded. After this, the electrode was cleaned with a dry cloth and set back at the same point for a new measurement to analyse repeatability.

We measured, grey, subcortical white matter, and white matter lying deep inside the brain 10 times at the same location. After this, the location was changed, and the measurement was repeated another 10 times.

We aimed to replicate the conditions of a real measurement situation during a surgery. That involves changes in the surface pressure each time the electrode is set on the tissue as well as changes of the measurement location.

### 2.5 Post processing

We averaged the pulses over 5 periods for noise suppression. In Fig. 4, four different current strengths of a 200 µs current pulse and their corresponding voltage responses in grey matter are shown.

**Figure 4:**
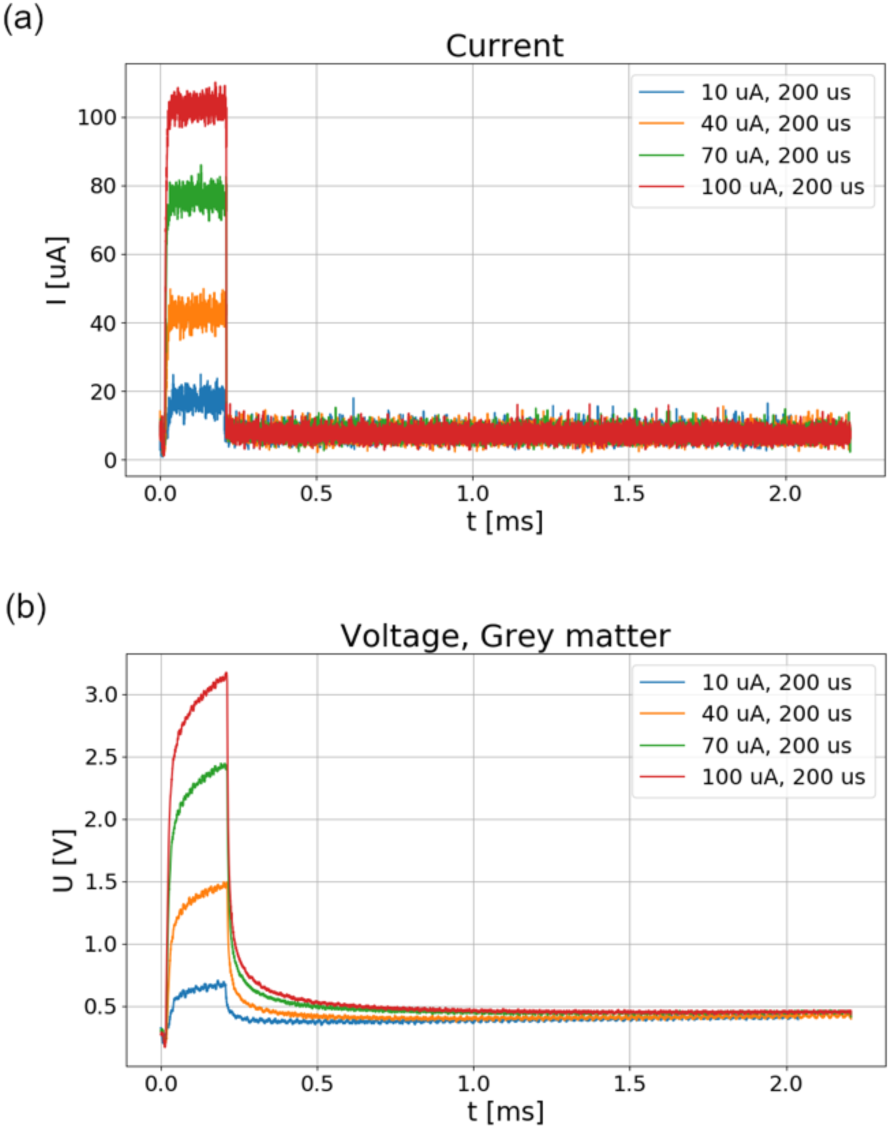
(a) Current pulses, current strengths: 10 µA, 40 µA, 70 µA, 100 µA, pulse width: 200 µs; (b) Measured voltages in grey matter, corresponding to current pulses above.

To compute the impedance, current and voltage were transformed to the frequency domain using the Discrete Fourier Transform (DFT):

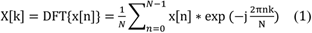

To obtain a higher frequency resolution, we zero-padded the voltage and current signals and doubled their length. This yielded a frequency resolution of 226.6 Hz.

Our approach, using time domain-based impedance detection, is an alternative to the use of sinusoidal measurement signals [7]. It yields a considerably reduced measurement time, which is very advantageous with regard to intraoperative measurements on humans.

The signals were sampled at 10 MHz. In theory, we could compute the impedance for up to 5 MHz. The stimulation signal quality, the electrode and its cable as well as the used measurement hardware limits the frequency range considerably. We set the limit to 200 kHz, mostly due to the spectrum of the current pulses.

The impedance was calculated as the quotient of the DFT of voltage and current:

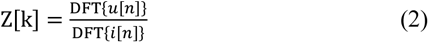

The DFT of a current pulse with a current strength of 100 µA and a pulse width of 200 µs as well as the DFT of the corresponding voltage for a measurement in grey matter are shown in Fig. 5. Note, that the amplitude at 200 kHz has already decreased by approximately 2 decades.

**Figure 5:**
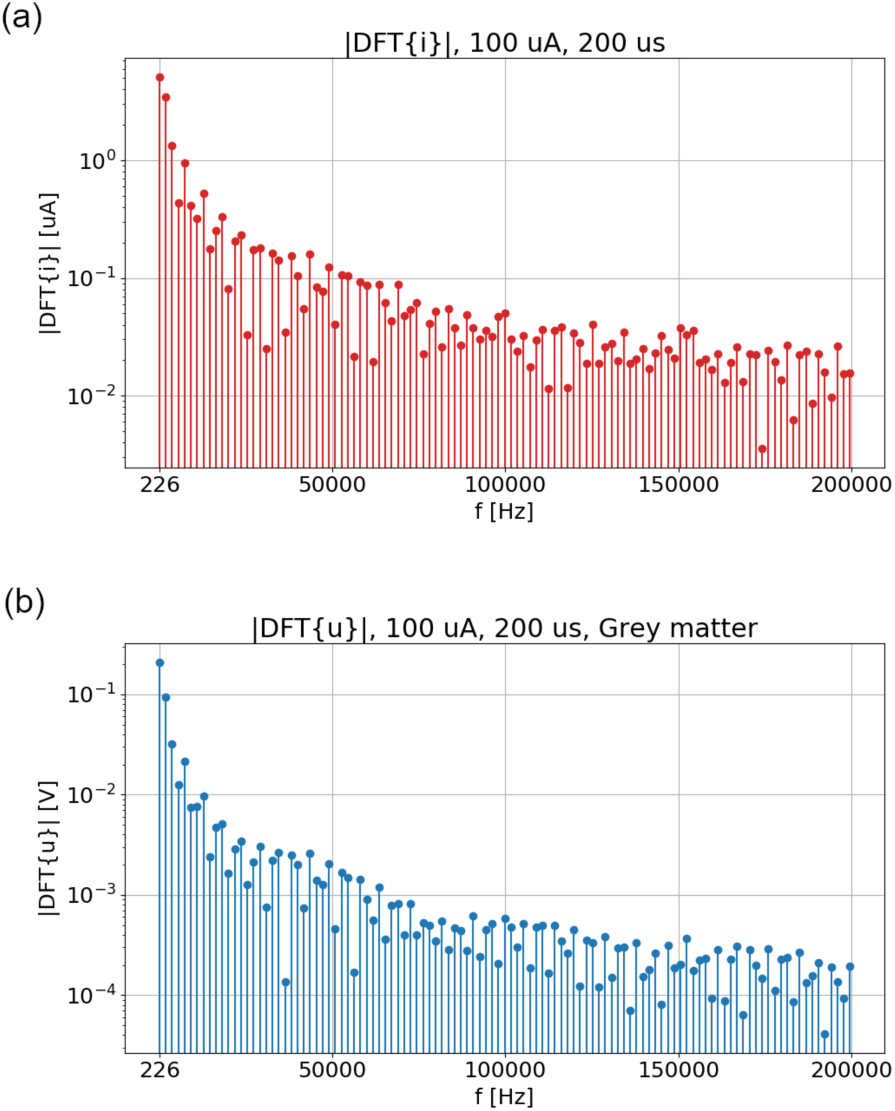
(a) DFT of current pulse, pulse strength: 100 µA, pulse width: 200 µs; (b) DFT of measured voltage response in grey matter.

The calculated impedance is rather noisy in the higher frequency range. To overcome this issue, the frequency axis was scaled logarithmically and values between two frequency points were averaged. This technique is known as logarithmic averaging and is mathematically correct, due to the usually dominating stochastic noise [7].

## 3 Results

The result of the current pulse of 100 µA and 200 µs meets the necessary requirements regarding signal-to-noise ratio and yields the most significant distinctness of the tissues. The impedance was computed for the 10 measurements on each tissue spot. In Fig. 6 the impedances of grey, subcortical white and deep white matter are shown. The plot is annotated with the p-values of a t-test between 10 measurements of grey and subcortical white matter. In the lower frequency range, the test is significant, while the p-values increase in the higher frequency range (bold annotation). The arrows indicate the span of the obtained results for 10 repetitions and the line represents the median.

**Figure 6:**
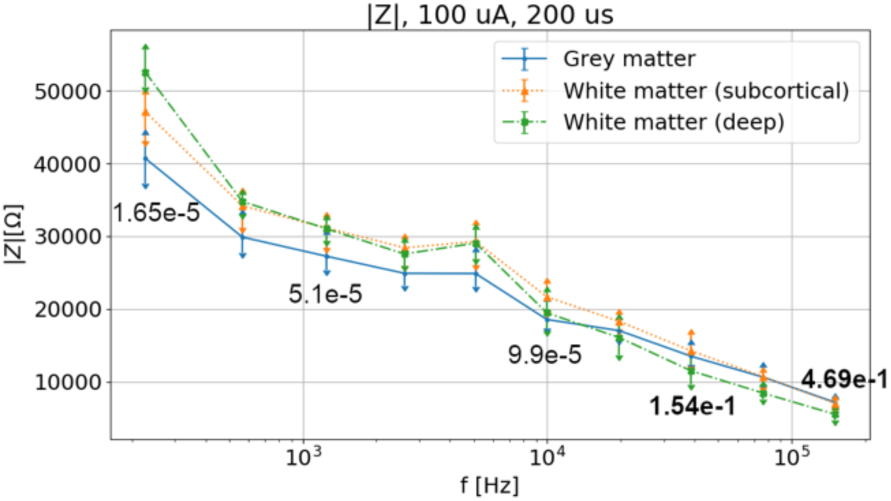
Median and span of computed impedances of grey matter, subcortical white matter, and deep white matter, pulse strength: 100 µA, pulse width: 200 µs.

## 4 Discussion

We could show the distinctness of grey and white matter with our measurement technique. It is noted that our results represent raw data, i.e. the total impedances, which are not yet corrected for the nonlinear electrode impedance. Current investigations are devoted to correct the collected impedance data for the individual electrode impedance. Therefore, a nonlinear electrode model will be developed and investigated by an extensive uncertainty and sensitivity analysis. Realistic head models and numerical simulations will be used to determine the electric field distributions during the measurements. Conductivity and permittivity can then be computed using the tissue impedance and the field distributions.

